# A Key Cytoskeletal Regulator of Ubiquitination Amplifies TGFβ Signaling During Mouse Developmental Vascular Patterning

**DOI:** 10.1101/055129

**Authors:** Ronak Shetty, Divyesh Joshi, Mamta Jain, Madavan Vasudevan, Jasper Chrysolite Paul, Ganesh Bhat, Poulomi Banerjee, Takaya Abe, Hiroshi Kiyonari, K. Vijayraghavan, Maneesha S. Inamdar

## Abstract

Vascular development involves *de novo* formation of a capillary plexus, which is then pruned and remodeled by angiogenic events. Cytoskeletal remodeling and directional endothelial migration are essential for developmental and pathological angiogenesis. Smad-dependent TGFβ signaling controls vascular patterning and is negatively regulated by microtubules. Here we show that a positive regulator of TGFβ signaling is essential for developmental vascular patterning and microtubule stability. Rudhira/BCAS3 is known to bind microtubules and to play a nodal role in cytoskeletal remodeling and directional endothelial cell (EC) migration in *vitro*. We demonstrate that the molecular and cellular function of Rudhira is deployed at critical steps in vascular patterning. We generated the first floxed mice for *rudhira* and find that global or endothelial knockout of *rudhira* results in mid-gestation lethality due to aberrant embryonic and extra-embryonic vessel patterning and defective cardiac morphogenesis. *Rudhira* null yolk sac ECs show random and retarded migration. Yolk sac transcriptome analysis revealed key mediators of angiogenic processes and TGFβ receptor signaling were perturbed in *rudhira* null mutants. Molecular and biochemical analyses showed that *rudhira* depletion reduced microtubule stability but increased expression of pathway inhibitors leading to high levels of SMAD2/3 ubiquitination and reduced activation. These effects were not rescued by exogenous TGFβ. However, TGFβ treatment of wild type ECs increased Rudhira expression. Further, exogenous Rudhira, which promotes directional cell migration, caused increased SMAD2/3 nuclear translocation and reduced inhibitor levels. Therefore, we propose that Rudhira and TGFβ signaling are mutually dependent. Rudhira has a dual function in promoting TGFβ signaling, possibly by sequestering microtubules and simultaneously preventing SMAD2/3 ubiquitination to permit EC migration and vascular patterning. TGFβ signaling and aberrant human Rudhira (Breast Cancer Amplified Sequence 3, BCAS3) expression are both associated with tumour metastasis. Our study identifies a cytoskeletal, cell type-specific modulator of TGFβ signaling important in development and cancer.

## Author Summary

Remodeling and fine patterning of the blood vasculature requires controlled and co-ordinated endothelial cell (EC) migration. This is achieved by the tight regulation of complex signaling pathways in developmental and adult vascular patterning. The TGFβ pathway, in particular, is important for this process. Here we show that the cytoskeletal protein Rudhira plays a critical role in promoting TGFβ signaling in EC. We generated “knockout” mice and show that Rudhira is crucial for EC migration in mouse developmental vascular patterning. In the absence of Rudhira there is increased degradation of TGFβ pathway effectors resulting in reduced signaling. This affects target gene expression and angiogenic processes, especially extracellular matrix remodeling and EC migration. As a result, the hierarchical vascular pattern is not achieved and embryos die mid-gestation. We propose that Rudhira plays a pivotal role in amplifying TGFβ signaling, which is critical for vascular remodeling in multiple normal and pathological contexts.

## Introduction

Vertebrate blood vessel formation involves *de novo* differentiation of endothelial cells (ECs) to form a primary plexus (1). This is resolved into a branched hierarchical network by EC migration and inhibition of proliferation, basement membrane reconstitution and recruitment of smooth muscle cells and pericytes to stabilize the network (2–4). Perturbation of the balance between pro- and anti-angiogenic cues leads to endothelial activation and cytoskeletal changes resulting in sprouting, migration and maturation (5, 6). This process is spatially and temporally orchestrated during development and adult neo–angiogenesis. Recent studies have elegantly elucidated the cellular dynamics of vessel regression and pruning (7, 8). EC migration and tube formation are key steps during angiogenesis, however, molecular mechanisms that operate are incompletely defined. Random and retarded EC migration results in an unpatterned, often leaky vasculature (9, 10).

A primary response of EC to angiogenic stimuli, flow induced shear stress or signals that induce pruning, is the reorganization of the cytoskeleton, resulting in activation of various signaling pathways (11–13). The loss of EC polarity, actin reorganization and extracellular matrix remodeling are key outcomes of cytoskeletal remodeling, resulting in directional cell migration essential for angiogenesis. The cytoskeleton also induces changes in cell shape and gene expression through a complex network of signaling pathways by influencing transcription factor activity (14).

The TGFβ pathway is sensitive to cytoskeletal rearrangements and regulates EC proliferation, differentiation, survival and migration (15). Multiple components of this pathway crosstalk with other signaling pathways to collaboratively pattern the vasculature (16). Mouse knockouts of TGFβ pathway components have demonstrated that a functional TGFβ pathway is essential for cardio–vascular development. Knockout of many of the ligands, receptors, effectors or regulators results in vascular developmental abnormalities and often causes embryonic death (17). In the embryonic yolk sac, paracrine TGFβ signaling does not affect EC specification or differentiation but regulates gene expression for assembly of robust vessels (18). Microtubules bind to and negatively regulate Smad activity (19). There is limited understanding of how cytoskeletal elements regulate the TGFβ and other signaling pathways and bring about changes in EC shape, polarity and directed migration. While several mouse mutant models of vascular patterning defects have been reported, those that perturb the cytoskeleton and associated proteins show pre-gastrulation lethality on one hand or display mild developmental phenotypes on the other (20, 21).

Here we investigate the role of an endothelial cytoskeletal protein Rudhira/BCAS3 in regulating developmental vascular remodeling. Rudhira is a WD40 domain-containing microtubule-binding protein expressed in vascular endothelial cells during mouse embryonic development (22). Rudhira promotes directional EC migration *in vitro* by rapidly re-localizing to the leading edge and mediating Cdc42 activation and filopodial extension (23).

We report that global or EC-specific *rudhira* deletion results in mid-gestation lethality due to aberrant vascular patterning. Knockout ECs fail to show directed migration. In *rudhira* mutants, hierarchical patterning of the embryonic and extra-embryonic vasculature is lost. Expectedly, this has a major effect on gene expression as seen by yolk sac transcriptome and protein expression studies. While several processes and signaling pathways involved in vascular patterning are perturbed, extracellular matrix remodeling and TGFβ signaling are majorly affected. We show that Rudhira does not induce, but promotes TGFβ signaling by suppressing expression of the inhibitor *smurf2,* thereby preventing SMAD2/3 ubiquitination. Further, Rudhira shows increased association with microtubules upon TGFβ induction. Thus Rudhira provides cytoskeletal control of several angiogenic processes by modulating TGFβ signaling. This indicates that normal Rudhira function is required to aid the EC cytoskeleton in regulating gene expression and cell migration. The identification and mechanistic deciphering of genessuch as *rudhira,* provide us a pivot on which molecular partnerships with both ubiquitous ‘essential’ or ‘redundant’ players can be understood and deciphered in the EC context.

## Results

### Endothelial *rudhira* is vital for embryonic development

Rudhira is expressed primarily in early embryonic vascular development and neo-angiogenesis, but its role *in vivo* is not known. Hence we generated *rudhira* floxed mice (Fig. 1A and S1A Fig.) and crossed them to *CMV-Cre* for ubiquitous deletion *(rudhira^flox/flox^; CMVCre^+^* abbreviated to *rudh^−/−^)* or *Tie–2–Cre* for tissue–specific ablation *(rudhira^flox/flox^; TekCre+* abbreviated to rudh^CKO^) of the *rudhira* locus (see Materials and Methods, and S1 Fig.). While heterozygotes were viable, ubiquitous or endothelial deletion of *rudhira* gave no live homozygous pups (S1A and S1B Tables). This indicates that endothelial deletion of *rudhira* causes recessive embryonic lethality. Analysis of embryos from E8.5 to E11.5 showed a reduced number of homozygous mutant embryos (S1A and S1B Tables) as identified by genotyping (Fig. 1B), transcript (Fig. 1C) and protein (S1B and S1C Fig.) expression, suggesting that lethality occurred between E9.0 and E11.5.

Since *rudhira* expression may be transient or undetectable in some migrating cells we also analysed the effect of globally deleted *rudhira* (*rudh*^−/−^) on development from E7.5 onwards, a stage before *rudhira* expression is initiated (22). At E7.5 *rudhira* mutant embryos were indistinguishable from controls with respect to morphology as well as primitive streak formation as seen by Brachyury expression (S1D Fig.). E8.5 mutant embryos showed unpatterned dorsal aorta as detected by Flk1 staining (S1E Fig.). By E9.5, mutant embryos were growth retarded (Fig. 1D and 2A) with defects including reduced somite number (19±2 at E9.5 in mutants compared to 25±2 in controls; n= 10) but expressed lineage markers of ectoderm (Nestin), mesoderm (Brachyury) and endoderm (AFP) (S1F Fig.). Rudhira is known to have restricted expression during vasculogenesis and primitive erythropoiesis (22). However, we did not find any significant change in the expression of vascular (CD34, Flk1, Ephrin, SMA, PECAM) and early hematopoietic (β-globin, c-kit, GATA-1) markers by semi-quantitative RT-PCR (S1F Fig.) showing that the two lineages are specified in *rudhira* null embryos. This suggests that *rudhira* is not essential for lineage specification and early vascular differentiation. Hence, we reasoned that the growth retardation and morphological defects seen in *rudhira* null embryos likely arise from defects in vascular rearrangement and patterning.

### Rudhira functions in extraembryonic vascular development

Impaired development and embryonic lethality between E8.5-E11.5 is often the result of aberrant and functionally impaired extra-embryonic vasculature (24). Moreover, Rudhira is strongly expressed in the yolk sac vasculature (22). Hence we analysed extraembryonic structures of mutant embryos, such as yolk sac and placenta which connect the maternal and fetal vasculature. Mutant yolk sacs were pale and lacked major blood vessels (Fig. 1D). Immunostaining for the blood vessel marker PECAM showed that *rudh*^−/−^ yolk sac vessels were irregular and fused, unlike the finely patterned honey-comb like vascular network seen in control littermates (Fig. 1E, 1I-K). Thus mutants could form a primitive vascular plexus which, however, did not undergo angiogenic remodeling. Primitive erythrocytes were dispersed all over the mutant yolk sac indicating an unpatterned and leaky vasculature. Histological analyses of yolk sac showed congested capillaries lined by thinner endothelium (Fig. 1F, arrowhead) as compared to controls (Fig. 1F, arrow). To test whether the yolk sac vessel patterning requires endothelial *rudhira* or is due to non-specific effects we analysed endothelial-deleted rudh^CKO^ yolk sac. Although of less severity than in the global knock-out, endothelial deletion of *rudhira* resulted in reduced branching from major vessels, vessel fusion and loss of branch hierarchy at both E10.5 and E11.5 (Fig. 1G–H and I–K). These results indicate that *rudhira* is essential for remodeling the yolk sac vascular network. Aberrant vascular remodeling in *rudhira* mutant embryos is likely the primary cause of death.

**Fig. 1.**
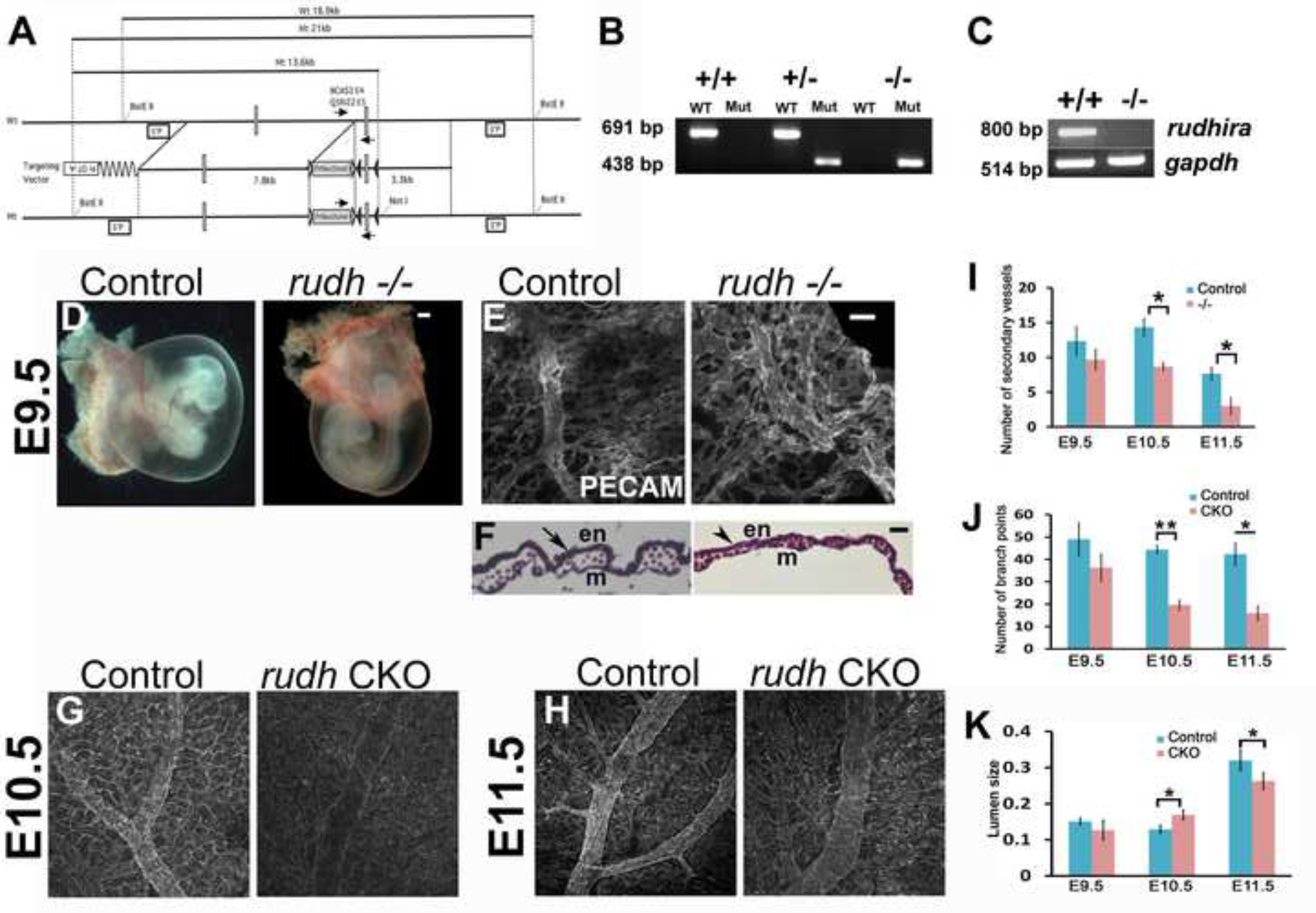
Rudhira is essential for vascular patterning in extraembryonic tissues. (A) Schematic showing strategy for generation of floxed allele of *rudhira* at exon 6. White rectangles indicate exons of the *rudhira* locus; black and white triangles indicate *loxP* and *frt* sequences respectively. 5′P and 3′P represent the probes that were used for Southern blot analyses. Arrows depict the positions of primers for routine genotyping of mice and embryos. (B) Genomic DNA PCR analyses showing genotype of control (+/+), heterozygous knock-out (+/−) and homozygous knock-out (−/−) embryos. (C) RT-PCR analysis showing *rudhira* mRNA expression in control (+/+) and homozygous knock-out (−/−) embryos. *GAPDH* RNA was used as a loading control. (D-H) Yolk sac morphology of control and *rudh−/− (rudhfl/fl;CMVCre+)* at E9.5 in whole mount unstained (D), immunostained for PECAM/CD31 (E) and sectioned and stained with hematoxylin-eosin (F). en: endoderm, m: mesoderm. (G, H) Yolk sac vasculature marked by PECAM staining in control and *rudh^CKO^ (rudhfl/fl;TekCre+)* at E10.5 and E11.5. (IK) Graphs showing quantitation of number of secondary vessels, branch points and lumen size in control and *rudh−/− (rudhfl/fl;CMVCre+)* at E9.5 and control and rudh^CKO^ *(rudhfl/fl;TekCre+)* at E10.5 and E11.5.

Placental circulation is vital for nourishment and development of the embryo. Improper development of the labyrinth, the feto-maternal interface, results in poor fetal invasion and causes growth retardation (25). Morphological analyses showed that *rudhira* null embryos have a smaller placenta (S2A, S2B and S2M Fig.) and with abnormal histology (S2A and S2B Fig.) as compared to controls of the same stage. Control placenta showed a distinct chorionic plate, labyrinth, spongiotrophoblast and decidual layers. *Rudh*^−/−^ placenta lacked stratified layers with a greatly reduced labyrinth and chorionic plate composed mostly of trophoblast giant cells. Fetal blood vessels could not invade into the placenta of *rudh*^−/−^ and contained fewer Ly76+ fetal erythrocytes as compared to controls where maternal (arrows) and fetal (arrowheads) blood cell pockets co-existed (S2C and S2D Fig.). Upon endothelial deletion of *rudhira* with *Tie2-Cre (rudh^CKO^),* placental thickness was reduced in mutant embryos at both E10.5 and E11.5 (S2E, S2F, S2I, S2J and S2M Fig.). Fetal vessel invasion was comparable to control at E10.5 (S2G and S2H Fig.) and was found to be affected only at E11.5 (S2K and S2L Fig.) although not as severely as in the global knock-out. This could explain the absence of significant growth retardation in conditional mutant embryos.

Taken together, these findings suggest that *rudhira* is essential for fetal vessel invasion into the developing placenta. Further, growth retardation in *rudh*“^/-^ embryos is likely a result of defective placental circulation.

### Rudhira plays a key role in cardiovascular development and tissue patterning

Since *rudh*^−/−^ embryos survived to E9.5 and *rudh^CKO^* survived to E11.5, we analyzed both genotypes for embryonic development and vascular patterning (Fig. 2A). Whole mount immunostaining of *rudh*^−/−^ embryos with anti–PECAM1 antibodies showed striking defects in the morphology and vasculature of the head, heart and intersomitic vessels (ISVs) (S3 Fig.). While control embryos had a well formed vascular network comprising major vessels giving rise to intricate secondary and tertiary branches (S3 Fig. arrows), *rudhira* mutants showed completely disorganized head vasculature with defective vessel sprouting, reduced capillaries and impaired branching of intersomitic vessels (ISV) that failed to sprout into fine capillaries (S3 Fig. arrowheads). Histological analysis as well as immunostaining for cardiovascular markers showed that both *rudh^−/−^* and *rudh^CKO^* embryos had collapsed, smaller heart chambers, reduced endocardium development and a fused atrio-ventricular canal. Dorsal aorta was discontinuous with a pronounced decrease in the lumen and intersomitic vessels were improperly patterned (Fig. 2B–D). The endothelial lining was disorganized in all tissues and ECs seemed to have impaired or random migration and were unable to form organized vessels. Similar phenotypes were observed in rudh^CKO^ at E10.5 and E11.5 (Fig. 2B–D).

**Fig. 2.**
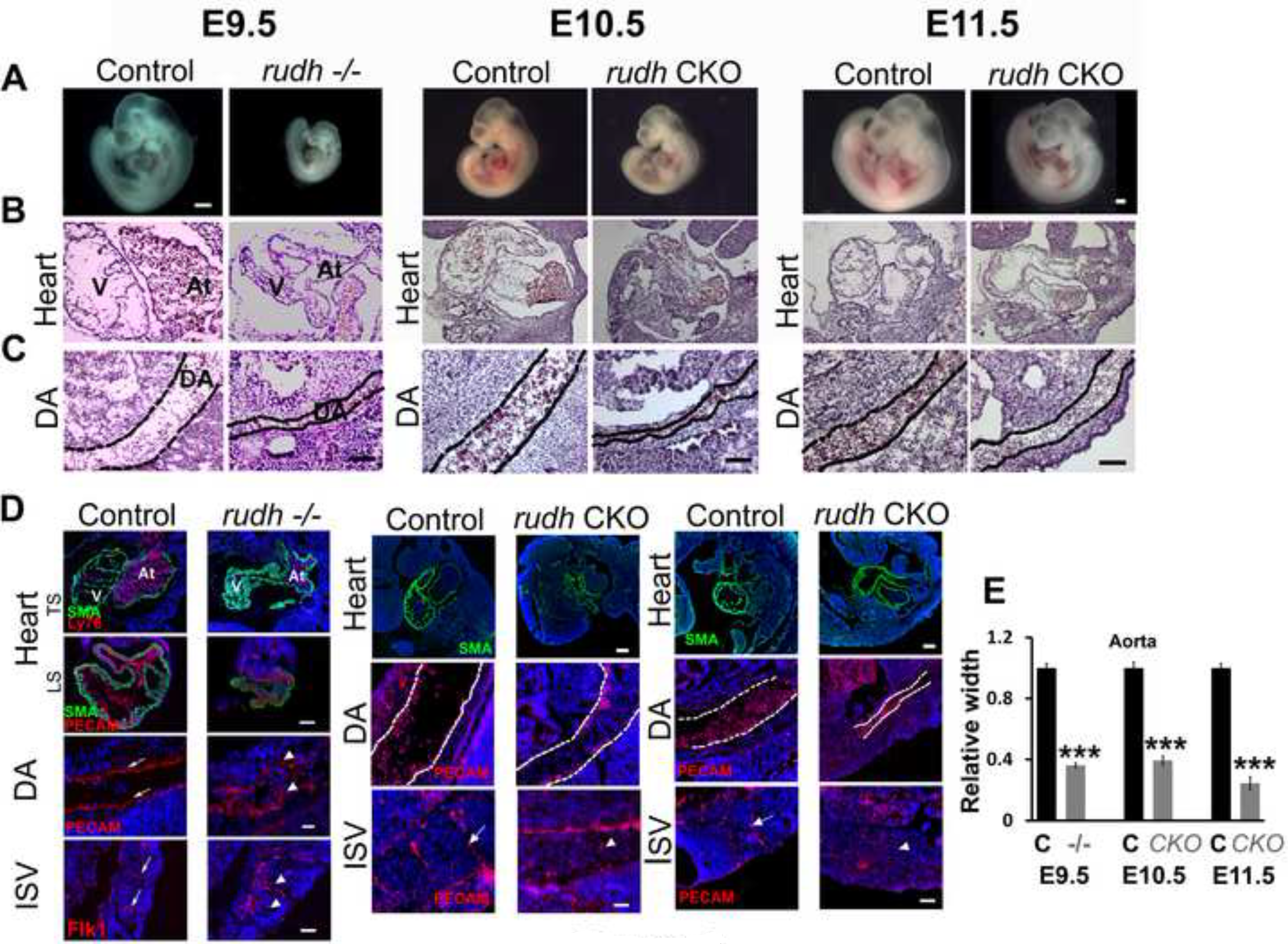
Absence of Rudhira leads to cardiovascular defects. Control and *rudh−/−* embryos at E9.5 or control and rudh^CKO^ at E10.5 and E11.5 were analyzed as indicated. (A) Unstained embryos, (B, C) Histological analysis showing comparison of heart and dorsal aorta (DA). (D) Immunostaining analysis of heart, dorsal aorta (DA) and intersomitic vessels (ISV) using myocardial marker SMA, primitive erythroid marker Ly76 and vascular markers PECAM and Flk1. Arrows indicate the normal vascular patterning and arrowheads point to irregular and discontinuous vasculature. TS: Transverse section, LS: Lateral section, V: Ventricle, At: Atrium. Nuclei are marked by DAPI (Blue). Black or white dotted lines mark the boundary of DA. (E) Graph shows quantitation of dorsal aorta width in the thoracic region. Results shown are a representative of at least three independent experiments with at least three biological replicates. Scale bar: (A) 500 μm; (B, C) 100 μm; (D) E9.5 heart: 100 μm, E10.5 and E11.5 heart: 200 μm, E9.5 DA: 20 μm, E10.5 and E11.5 DA: 50 μm, ISV: 50 μm.

To test whether *rudhira* null ECs also show slow and random migration, we cultured yolk sac ECs and tested them in a wounding assay (Fig. 3A). *Rudhira* null ECs also showed reduced migration rate and decrease in directed migration (Fig. 3B and 3C). Taken together, our results demonstrate a key role for Rudhira in directed cell migration, essential for vascular remodeling during developmental angiogenesis.

**Fig. 3.**
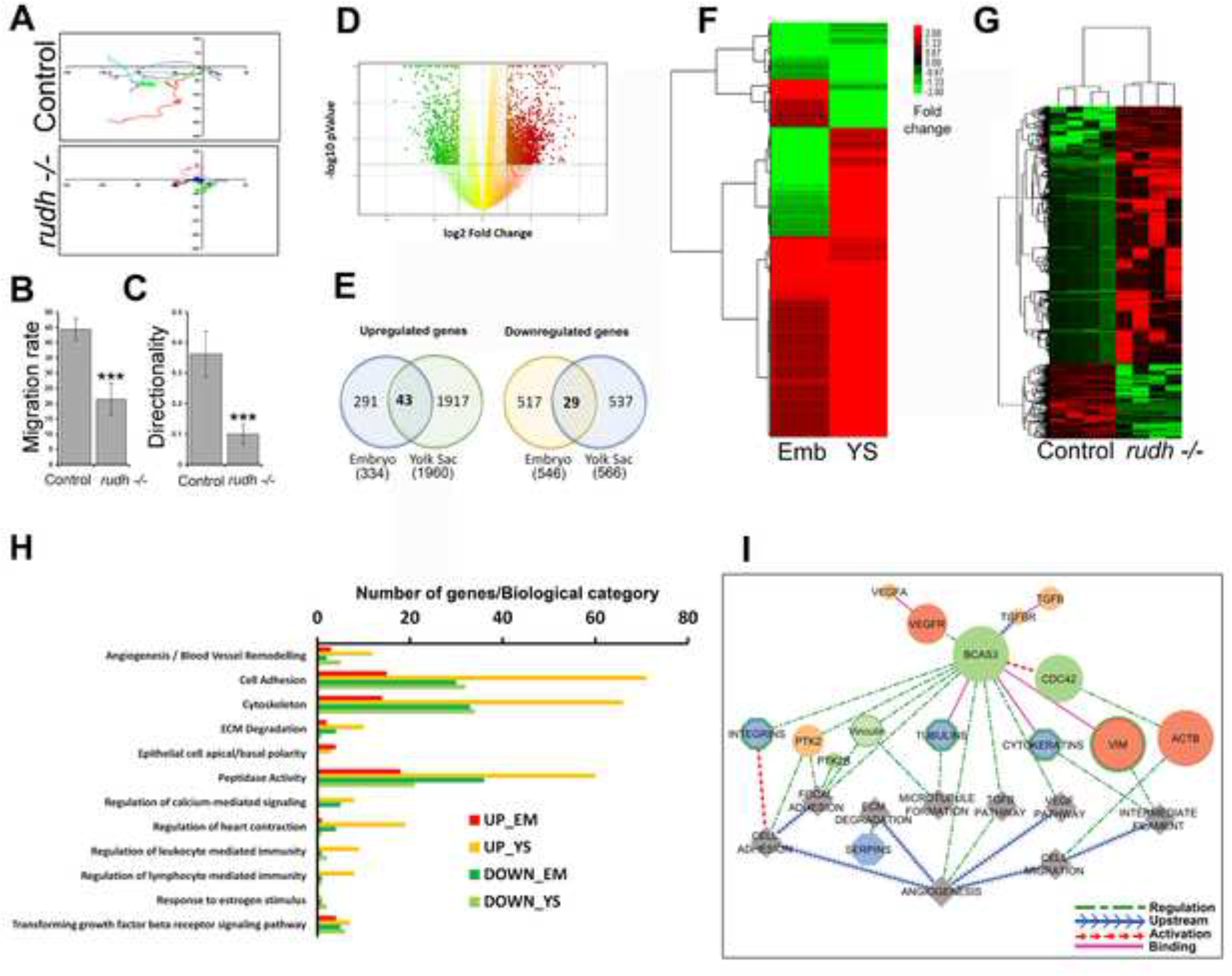
Rudhira depletion causes impaired cell migration and promotes yolk sac transcriptome expression. (A-C) Migration tracks (A) of control and *rudh^−/−^* yolk sac endothelial cells subjected to wounding assay. Quantification of the rate of migration (B) and directionality (C) compared between control and *rudh^−/−^* yolk sac endothelial cells. At least 30 cells were analysed per genotype. (D) Representation of differentially expressed genes in *rudh^−/−^* by volcano plot. (E-H) Transcriptome analysis. (E) Venn diagram showing number of genes (unique probe sets) dysregulated in embryo and yolk sac upon *rudhira* deletion. (F) Unsupervised hierarchical clustering of differentially expressed gene changes in *rudh^−/−^* embryo (Emb) and yolk sac (YS) at E9.5 compared to controls. Each row represents a gene. Expression intensities are displayed from green (low expression) to red (high expression). Lines on the left represent the similarity between genes with the most similar expression profiles clustered together with the shortest branches and represented in the dendrogram to illustrate their relationship. (G) Unsupervised hierarchical clustering of differentially expressed gene changes in *rudh^−/−^* yolk sac at E9.5 compared to controls. Each row represents a gene, and column represents the tissue. (H) Histogram showing significantly enriched Gene Ontology (GO) and Pathways (p<=0.05) harboring differentially expressed genes in the embryo and yolk sac upon *rudhira* deletion. (I) Model depicting *rudhira/BCAS3* gene regulatory pathway. Cytoscape V 8.0 was used to visualize the network. Results shown are a representative of two biological and two technical replicates for microarray and at least three independent experiments with at least three biological replicates for other experiments. Statistical analysis was carried out using one-way ANOVA. Error bars (in B and C) indicate mean ± SD. ^***^p<0.001.

### Rudhira regulates expression of the angiogenesis network

Rudhira regulates actin dynamics which is known to affect nuclear transcription. We performed whole transcriptome-based analysis of gene expression in *rudhira* knockout yolk sac and embryos at E9.5 to determine its effect on vascular remodeling. While Rudhira expression is primarily in the endothelium, our studies *in vitro* ((23) and this report) indicated that changes in Rudhira expression and localization are rapid and depend on inter-cellular and cell-ECM interactions. Hence we chose to analyze intact tissue with minimal manipulation to understand how Rudhira affects the transcriptome. While embryos at E9.5 have a diverse set of derivatives of all three germ layers, yolk sacs are primarily made of primitive endoderm and mesoderm, the latter comprising mainly endothelial and hematopoietic lineages. Hence we separately analyzed *rudh^−/−^* embryos and yolk sacs to avoid confounding effects of possible mosaic deletion of *rudhira* in the *Tie2-Cre* conditional knockout. We then extensively validated the data by quantitative PCR-based expression analysis of yolk sac RNA and endothelial cell line RNA. Volcano plot based method was used to visualize the transcripts that are two-fold differentially expressed in yolk sac (Fig. 3D). 3291 unique probes showed 2-fold or greater statistically significant changes in gene expression (S2 Table). Of these 546 were downregulated and 334 upregulated in embryo and 566 downregulated and 1960 upregulated in yolk sac (Fig. 3E). 29 downregulated genes and 43 upregulated ones were common between embryo and yolk sac (Fig. 3E and S3 Table). Unsupervised hierarchical cluster analysis showed that genes with similar expression patterns were clustered together with branch distance proportional to their similarity in expression pattern. A distinct sub set showed reciprocal expression between embryo and yolk sac (Fig. 3F). Interestingly the majority of clustered genes were mainly upregulated in the yolk sac (Fig. 3F and 3G), while the embryo had a more balanced distribution in each cluster (Fig. 3F). To define how changes in gene expression caused by *rudhira* depletion may influence vascular development and remodeling, we functionally annotated the data using DAVID (Database for Annotation, Visualization and Integrated Discovery) and found that genes linked to many biological pathways were enriched. Key deregulated biological categories were identified (Fig. 3H).

Analysis of the entire data set showed greater variation between duplicates of embryo than yolk sac, possibly because of heterogeneity in the embryonic tissue. Hence for further analysis we focused on yolk sac data as it is also the primary site of vascular remodeling and shows early Rudhira expression. Significant expression changes were seen in genes that relate to a range of processes or pathways, which could impact on vascular development and remodeling (Fig. 3H and S4 Fig. and S5 Table). Gene ontology analysis of the common genes identified principal biological processes affected by the loss of *rudhira* with a Z score above 2.5. Amongst signaling pathways, negative regulation of the transforming growth factor beta (TGFβ) signaling pathway was identified as significantly perturbed. Other pathways implicated in vascular development such as Wnt, JAK/STAT and Notch signaling showed changed levels of a few genes but expression of the majority of the pathway components was unaffected. Important regulators of cellular processes such as angiogenesis/blood vessel remodeling, extracellular matrix, regulation of proteolysis, negative regulation of peptidase activity and cell projection organization were identified. Further, key molecular families involved in cytoskeletal remodeling, cell adhesion, cell migration and TGFβ and VEGF pathways (all important during angiogenesis) were connected by Rudhira/ BCAS3 allowing us to identify the Rudhira network in angiogenesis (Fig. 3I and S6 Table).

### Identification of regulatory networks and nodes regulated by Rudhira

A total of 140 genes from cluster analysis were enriched in the key gene ontology (GO) and pathways identified (S5 Table) with a significance criterion of p<0.05. Further, we were able to associate GOs and pathways known to co-operate during vascular development and remodeling namely adhesion, angiogenesis, cytoskeleton, ECM organisation, peptidase activity and TGFβ signaling (Fig. 3I). Genes differentially expressed in these six processes were subjected to unsupervised hierarchical clustering to identify molecular signatures (Fig. 4A, 5A and S4 Fig.). An interaction network of significant GO terms was assembled into a GO map to depict the relationship among prominent functional categories (Fig. 4B, 5B and S4 Fig.). Subsequent verification of expression data was carried out for key genes known to mediate these processes (Fig. 4C, 5C, 5D and S4 Fig.). 51 out of the 3407 genes that showed significant variation from control in the knockout yolk sac were validated by qPCR on cDNA generated from *rudhira* knockdown and non-silencing control endothelial cell line RNA (Fig. 4C, 5C, 5D and S4 Fig.) and 70% of these (36/51) agreed with array data. Changes in expression level of selected candidates were further validated using cDNA generated from fresh E9.5 yolk sac RNA (S4F Fig.).

**Fig. 4.**
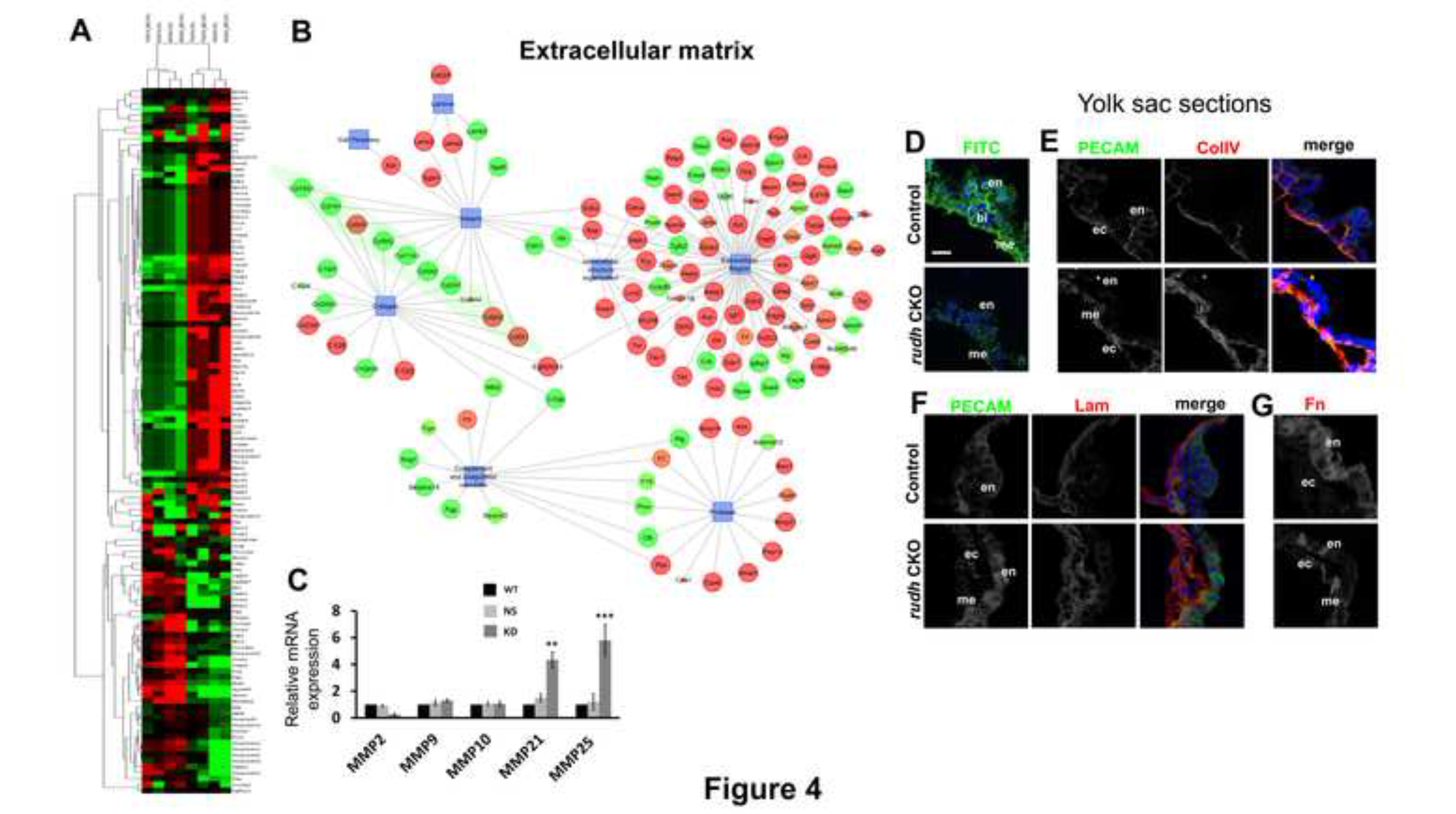
Rudhira depletion affects extra cellular matrix (ECM) organization around blood vessels. (A) Unsupervised hierarchical clustering and (B) regulatory network of the extracellular matrix (ECM) organization biological process showing genes significantly deregulated in *rudh*^−/−^ yolk sac compared to control. Edge weighted spring embedded layout was used to visualize the network (see methods). (C) Graph shows relative transcript levels of selected *MMP* genes from the ECM network that were validated by qRT PCR on wild type (WT), non-silencing control (NS) and *rudhira* knockdown (KD) lines (SVEC). (D-G) Yolk sac sections from control and rudh^CKO^ at E10.5 showing (D) live *in situ* gelatin zymography (FITC, green), (E-G) co-expression of vascular marker PECAM with (E) CollagenIV (ColIV), (F) Laminin, (G) Fibronectin (Fn). Nuclei are marked by DAPI (Blue). Error bars indicate standard error of mean (SEM). Results shown are a representative of at least three independent experiments with at least three biological replicates. Statistical analysis was carried out using one-way ANOVA. Scale bar: (D-G) 20 lμm (see also S5 Fig.). ^**^p<0.01, ^***^p<0.001.

### Endothelial Rudhira regulates extracellular matrix (ECM) organization

Yolk sac transcriptome analysis revealed several processes important for angiogenesis that were perturbed upon *rudhira* depletion. Extracellular matrix was an over-represented functional category, in which genes for matrix components and ECM proteases are aberrantly expressed (Fig. 4A and 4B). Controlled ECM remodeling is important for cell migration. Mutations that affect the dynamic regulation and crosstalk between the cell and ECM are also known to affect tissue patterning and homeostasis (26). *MMP10, MMP21* and *MMP25* were aberrantly expressed while *TIMPs* (Tissue inhibitors of metalloproteases) were not significantly changed in *rudh^−/−^* yolk sacs. Expression of a large number of the serine protease inhibitors was aberrant with eight members downregulated and eleven upregulated (Fig. 4B and S2 Table). Interestingly several members of the extracellular *Serpina* clade were downregulated (6/8) whereas the intracellular *Serpinb* clade was upregulated (8/9). Serpinb clade members inhibit Granzyme (Gzm) activity (27). In the *rudh^−/−^* yolk sac *Gzmd* and *Gzmg* transcripts were upregulated suggesting a loss of balance between production of proteases and protease inhibitors. A disintegrin and metallopeptidase domain (Adam) class of endopeptidases namely *Adam11, Adam24, Adam32, Adam7, Adamts12, and Adamts15* were all significantly upregulated in *rudh^−/−^* yolk sacs. This suggested that matrix organization or degradation may be affected.

ECM degradation is an essential step in angiogenesis and vascular patterning (28, 29). Collagen type IV is a major component of the vascular basement membrane. Of 11 collagen family members affected by loss of *rudhira,* 7 were downregulated (Fig. 4B). *In situ* zymography on live *rudh^CKO^* yolk sac and embryo sections at E10.5 using dye quenched (DQ) gelatin showed absence of or significant decrease in signal around vessels indicating reduced matrix degradation as compared to controls (Fig. 4D and S5A Fig.). Sections were stained for collagen and laminin post zymography (S5A and S5D Fig.). Both *rudh^CKO^* yolk sacs and embryos showed highly disorganized collagen around the vessels (Fig. 4E and S5B Fig.). Further, Rudhira expression in vessels of control embryos overlapped with collagen staining (S5B and S5C Fig.). Co-staining of fixed sections of yolk sac and embryos with endothelial marker PECAM and ECM markers showed that the organization of the matrix components collagen, fibronectin and laminin was disrupted (Fig. 4E–G, S5 Fig.). These results indicate that Rudhira is essential for maintaining proper ECM organisation. TGFβ signaling plays a major role in regulating genes involved in ECM deposition and degradation. As levels of several transcripts of the TGFβ pathway were altered in *rudhira* mutant yolk sac (Fig 5A, 5B and 5H) we analysed this pathway further.

### Regulation of TGFβ signaling by Rudhira

TGFβ pathway was one of the major over-represented categories as levels of several of the TGFβ pathway molecules were altered in *rudhira* mutant yolk sac (Fig 5A, 5B and 5I). The TGFβ pathway is essential for vascular patterning and angiogenesis. Deletion of TGFβ1 leads to delayed wound healing (30). Ablation of TGFβRI or TGFβRII results in embryonic lethality with severe patterning defects (31, 32). A significant decrease in expression of *TGFb2, TGFb3* and *TGFbr2* was seen in the microarray analysis and subsequent validation by qRT-PCR (Fig 5A–D and Fig 5I). TGFβ antagonists like *Smurf1, Smurf2* and *Smad7* were upregulated in *rudhira* knockdown ECs (Fig 5D). TGFβ is produced by the yolk sac ECs and is also available as a paracrine signal from the yolk sac endoderm and embryonic tissue (18). This activates the SMAD2 effector which leads to the differentiation of vascular smooth muscles cells and promotes their association with endothelial cells, to establish a properly patterned vasculature. Reduced pSMAD2 levels in *Smad2/3* and *endoglin* mutants co-relate with an unpatterned vasculature and mid-gestation lethality (18, 33). To assess pSMAD2 levels in the absence of Rudhira, we stained control and *rudh^CKO^* yolk sacs for pSMAD2. Mutant yolk sacs had fewer pSMAD2 positive endothelial cells (Fig. 5E, arrows). Upon induction of live yolk sacs with exogenous TFGP, control yolk sacs showed increased pSMAD2, whereas *rudh^CKO^* yolk sacs continued to show low pSMAD2 signal in the endothelial cells (Fig. 5E, arrowheads). Similarly, *rudhira* knockout embryo-derived endothelial cells or *rudhira* knockdown endothelial cell line (SVEC KD) treated with TGFβ also failed to activate pSMAD2 as compared to controls (Fig. 5F and Fig. 5G respectively, and S6A Fig.). This suggests that Rudhira is essential for TGFβ pathway activation via pSMAD2.

**Fig. 5.**
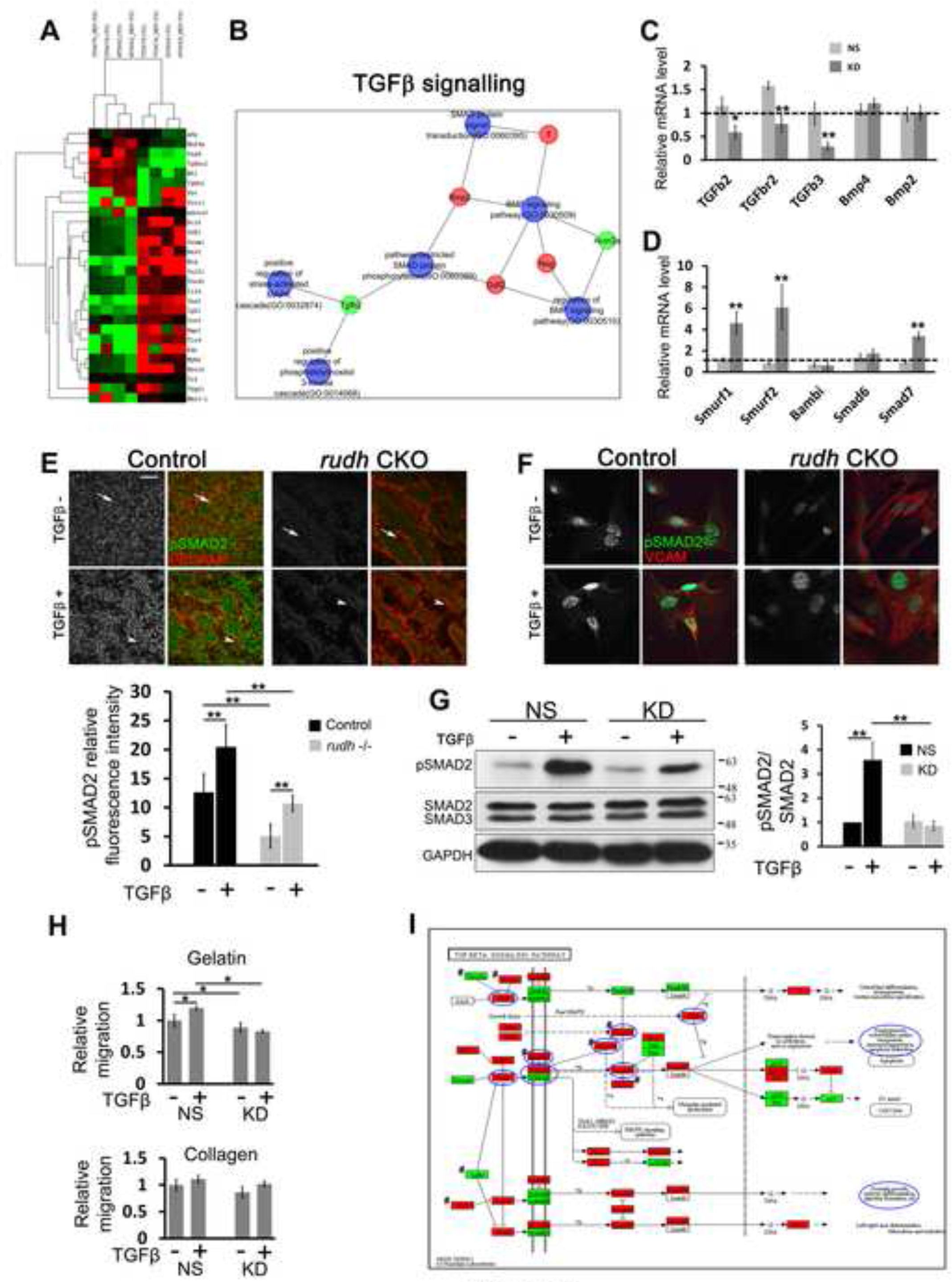
Rudhira depletion deregulates TGFβ signaling machinery essential for angiogenesis. (A, B) Unsupervised hierarchical clustering computed with Pearson Uncentered algorithm with average linkage rule and regulatory networks of differentially expressed genes and biological processes significantly deregulated in *rudh−/−* yolk sac compared to control. Network maker program (Bionivid Technology Pvt Ltd) was used to identify the nodes and edges that form the regulatory circuit and Cytoscape V 8.0 was used to visualize the network. Edge weighted spring embedded layout was used. (C-D) Graphs show representative positive regulators (C) and negative regulators (D) of the TGFβ pathway that were validated by qRT PCR on wild type (WT), non-silencing control (NS) and *rudhira* knockdown (KD) endothelial cell (EC) lines (SVEC). (E-F) Untreated and TGFβ treated control and rudh^CKO^ *(rudhfl/fl;TekCre+)* yolk sacs (E) or yolk sac derived cells (F) were analyzed for the expression of phosphorylated SMAD2 (pSMAD2) by immunostaining. Samples were co-stained with endothelial markers PECAM (E) or VCAM (F). Graphs show mean fluorescence intensity for pSMAD2. n = 3 yolk sacs. (G) Untreated and TGFβ treated non-silencing control (NS) or *rudhira* knockdown (KD) SVEC lines were analyzed for expression of TGFβ pathway molecules by immunoblot of cell lysates. Graph shows pSMAD2/SMAD2 ratio. (H) Graphs showing relative migration of control (NS) and *rudhira* knockdown (KD) SVEC lines plated on collagen or gelatin- coated dishes and induced with TGFβ. Error bars indicate standard error of mean (SEM). Results shown are a representative of at least three independent experiments with at least three biological replicates. Statistical analysis was carried out using one-way ANOVA. Scale bar: (E) 20 μm ^*^p<0.05, ^**^p<0.01, ^***^p<0.001. (I) KEGG TGFβ Pathway. Genes in the TGFβ pathway from KEGG database mapped based on fold change in *rudh−/−* yolk sac in comparison to control to understand pathway regulation. Green: downregulated, Red: upregulated. # indicates negative regulators and antagonists of TGFβ pathway. Blue circles indicate genes whose expression level was validated in this study or processes that were found to be affected.

### Rudhira functions downstream of TGFβ receptor activation

To test whether provision of TGFβ could rescue the cell migration defect seen upon *rudhira* depletion, we cultured SVEC knockdown cells in the presence of exogenous TGFβ. Provision of exogenous TGFβ in a cell monolayer wounding assay did not rescue the migratory defects of *rudhira-depleted* endothelial cells migrating on gelatin (Fig. 5H). This suggested that the TGFβ pathway requires Rudhira for activating processes that promote cell migration. Since loss of *rudhira* results in an inability to activate TGFβ- mediated Smad2 signaling, this could lead to an inability to migrate, resulting in angiogenic defects. Further, *rudhira* depletion cell autonomously affects TGFβ-dependent migration. Alternatively, since Rudhira affects multiple signaling pathways, TGFβ alone may not be sufficient to rescue the effect of *rudhira* deletion.

### Rudhira is required for microtubule stability

Microtubules (MT) bind to TGFβ pathway effectors Smad2 and Smad3 thereby preventing their phosphorylation and activation (19). Binding of Smads to MT is independent of TGFβ stimulation. TGFβ triggers dissociation of the Smad complex from MT and increases Smad2 phosphorylation and nuclear translocation. Our earlier work (23) showed that Rudhira interacts with MT, which prompted us to investigate whether Rudhira is required for MT-regulated TGFβ signaling. Rudhira depletion led to reduced alpha-tubulin as well as acetylated tubulin levels in SVEC endothelial cell line as well as *rudhira* null primary ECs (Fig. 6A,B). MTs containing acetylated and detyrosinated tubulin are known to re- localize towards the leading edge of a migrating cell (34). In a monolayer wounding assay Rudhira-depleted cells showed reduced acetylated tubulin staining, which was not localized towards the leading edge (Fig. 6C). Further, upon TGFβ stimulation, wild type ECs (SVEC) showed increased co-localization of Rudhira with MT, suggesting increased binding (Fig. 6D). This suggests a role for Rudhira in regulating MT architecture and MT- regulated TGFβ signaling.

### Rudhira augments TGFβ pathway activation

To test whether Rudhira alone is sufficient to activate the TGFβ pathway, we transfected endothelial cells (SVEC) with a Rudhira overexpression construct *(Rudh2AGFP)* and analyzed the status of TGFβ pathway activation. Upon ligand binding to the TGFβ receptor, Smad2/3 gets activated and translocates to the nucleus (35, 36). Rudhira overexpression alone was not sufficient to activate TGFβ signaling and cause Smad2/3 nuclear localization (Fig. 6E). However, upon addition of TGFβ1 to the culture, Rudhira overexpressing cells showed markedly increased pathway activation as seen by increased nuclear Smad2/3 in *Rud2AGFP* transfected cells compared to un-transfected or vector control cells. Further, Rudhira overexpression is unable to overcome the receptor-level inhibition of TGFβ pathway using the small molecule inhibitor SB431542 (SB) that binds to the Alk5 co-receptor, thereby blocking pathway activation (Fig. 6E). This confirms that Rudhira can augment pathway activity but not substitute for TGFβ receptor activation. It also indicates that Rudhira functions downstream of the TGFβ receptor. This is in agreement with the cytoskeletal localization of Rudhira.

**Fig. 6.**
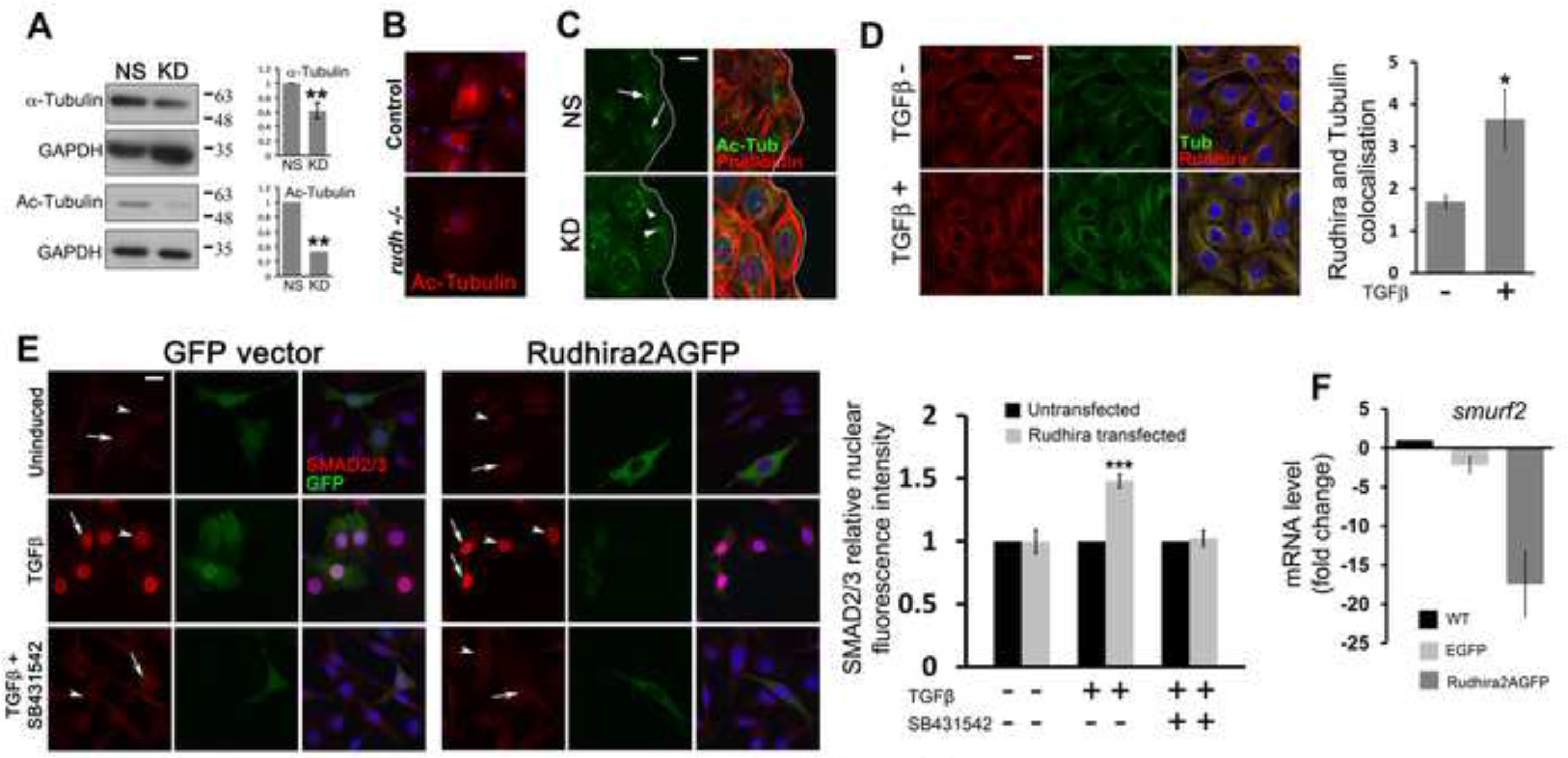
Rudhira stabilizes microtubules and augments TGFβ pathway activation. Non-silencing control (NS) or *rudhira* knockdown (KD) ECs were analysed for microtubule organization and stability. (A–C) Expression of α-Tubulin and acetylated-Tubulin (Ac-Tubulin) by immunoblot of cell lysates (A) and localisation of acetylated-tubulin by immunostaining of fixed yolk sac cells (B) and SVEC NS and KD cells (C). (D) Colocalisation analysis of Rudhira and microtubules on TGFβ induction. Graph shows quantitation of Rudhira and MT colocalisation from at least 20 cells. (E) SVEC were transfected with plasmid constructs for expression of GFP or Rudh2AGFP, and treated with either TGFβ or TGFβ+ SB431542 (TGFβ inhibitor) and analysed for SMAD2 by immunostaining. Graph shows quantitation of SMAD2/3 nuclear fluorescence intensity. Compare arrows (transfected cell) and arrowheads (untransfected cell). (F) qRT PCR analysis of *smurf2* transcript levels in wild type (WT), GFP or Rudhira2AGFP expressing endothelial cells (SVEC). Error bars indicate standard error of mean (SEM). Results shown are a representative of at least three independent experiments with at least three biological replicates taken into account. Statistical analysis was carried out using one-way ANOVA. Scale bar: (B–E) 20 μm. ^*^p<0.05, ^**^p<0.01, ^***^p<0.001.

Interestingly, Rudhira overexpression also led to a decrease in *smurf2* levels (Fig. 6F and S7C Fig.). This indicates that Rudhira augments TGFβ pathway activation by increasing SMAD2/3 activation, possibly due to a reduction in inhibitor levels. To test whether the converse is true, i.e. whether TGFβ affects Rudhira expression, we treated wild type SVEC cultures with TGFβ1. There was a significant increase in Rudhira transcript and protein levels upon TGFβ induction, as detected by qPCR and immunofluorescence staining (S7A, B Fig.). This indicates that Rudhira and the TGFβ pathway are mutually dependent for their expression and function.

### Rudhira inhibits Smurf- mediated TGFβ pathway attenuation

Our analysis indicated that Rudhira is not an inducer but a promoter of the TGFβ signaling pathway in endothelial cells that acts downstream of receptor-level activation but upstream to SMAD activation. KEGG pathway analysis of the yolk sac transcriptome showed that a majority of the genes involved in TGFβ signaling were deregulated upon *rudhira* loss. Especially, several antagonists *(noggin, follistatin, bambi)* and negative regulators *(sara, smurf1/2, smad6/7)* of the pathway were upregulated (Fig. 5I).

**Fig. 7.**
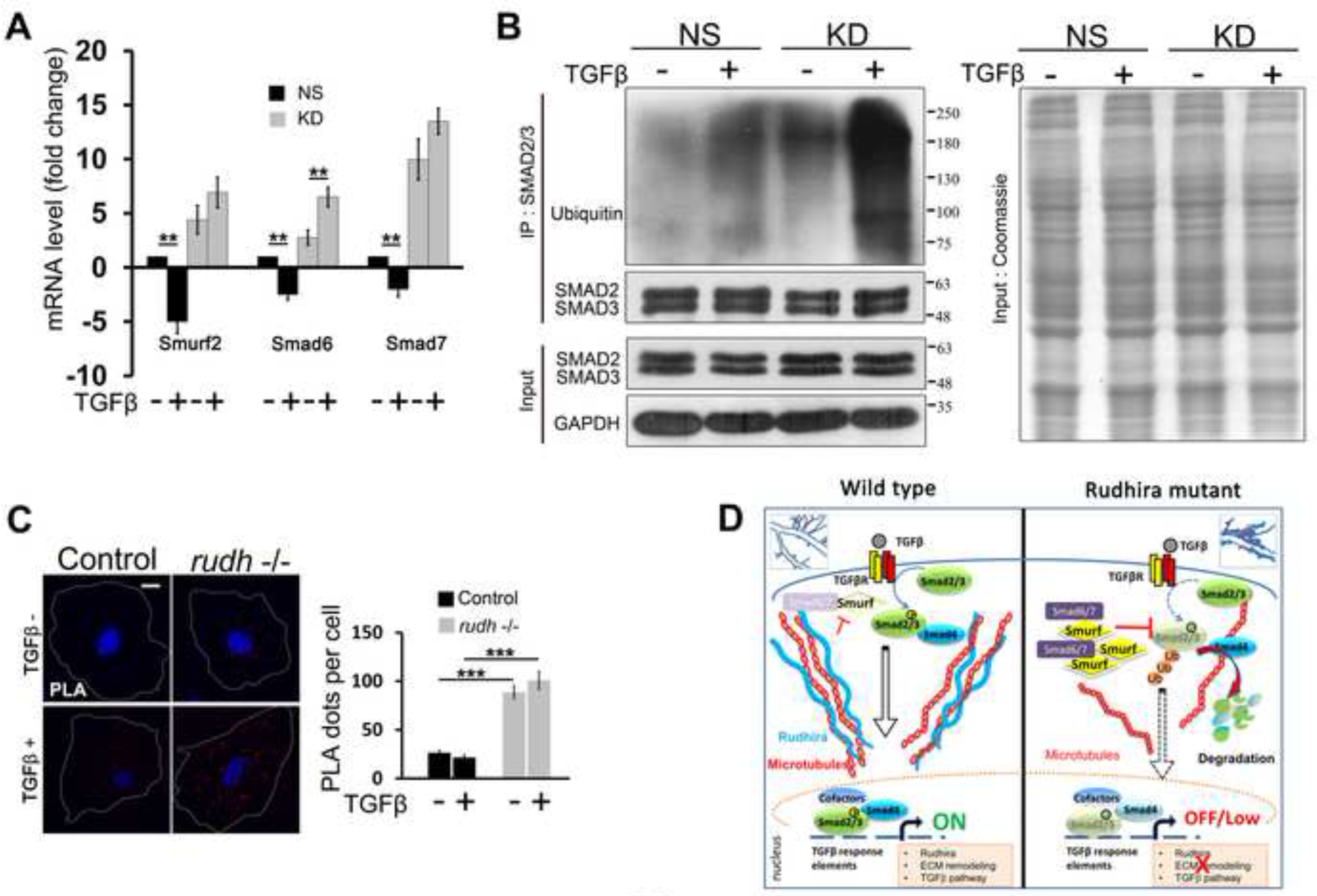
Rudhira inhibits Smurf- mediated inhibition of TGFβ pathway. (A) qRT PCR analysis of negative regulators of TGFβ pathway in non-silencing control (NS) and *rudhira* knockdown (KD) endothelial cell (EC) lines (SVEC) upon TGFβ treatment. (B) Untreated and TGFβ treated non-silencing control (NS) or *rudhira* knockdown (KD) SVEC lines were analysed for Smad2/3 ubiquitination by immunoprecipitation (IP) of Smad2/3 and immunoblotting for ubiquitin. Cell extracts were resolved by SDS-PAGE and either Coomassie- stained or immunoblotted for indicated proteins. (C) *in situ* Proximity ligation assay (PLA) for Smad2/3 and Ubiquitin on control and *rudh^−/−^* yolk sac cultured cells with or without TGFβ treatment. PLA dots represent ubiquitinated Smad2/3. Graph shows the quantitation of PLA dots per cell. (D) Schematic representation of the role of Rudhira in vascular patterning. The status of TGFβ signaling and microtubules in the presence (wild type) or absence (Rudhira mutant) of Rudhira is shown. Boxed insets show blood vessel (red) branching and ECM (blue lining) in wild type and *rudhira* mutant yolk sacs. Error bars (A-C) indicate standard error of mean (SEM). Results shown are a representative of at least two independent experiments with at least two biological replicates. Statistical analysis was carried out using one-way ANOVA. ^*^p<0.05, ^**^p<0.01, ^***^p<0.001.

The TGFβ pathway is regulated at multiple steps by a complex combination of activators, inducers and inhibitors (37). Ubiquitin- mediated proteasomal degradation is essential for fine regulation of TGFβ pathway activity in the basal as well as activated states. The E3 ubiquitin ligases for SMADs, namely Smurf1 and Smurf2 (SMAD specific E3 Ubiquitin ligases) are important regulators of the pathway. They control SMAD2/3 levels by ubiquitination and targeting for proteasomal degradation (38, 39). These inhibitors are also targets of TGFβ pathway activation.

Since Rudhira depletion results in increased *Smurf1/2* and inhibitory Smad *(Smad6* and *Smad7)* levels, we checked the effect of exogenous TGFβ addition on these inhibitors. While control cells show a sharp decrease in the levels of these inhibitors upon addition of TGFβ, Rudhira- depleted cells showed no reduction in their levels, suggesting a loss of control on the negative regulators of TGFβ signaling (Fig. 7A). To test whether increased inhibitor transcript levels co-relate with function, we assayed for levels of ubiquitinated-SMAD2/3 by immuno-pulldown of SMAD2/3 followed by immunoblotting for Ubiquitin.

Rudhira depletion indeed results in increased SMAD2/3 ubiquitination, which is markedly increased on TGFβ- mediated activation of the pathway (Fig. 7B). To further check the status of SMAD2/3 ubiquitination *in vivo,* we performed a Proximity Ligation Assay (PLA) for SMAD2/3 and Ubiquitin in primary endothelial cell cultures derived from E9.5 control and *rudh−/−* yolk sacs. PLA is a robust and sensitive assay to identify post-translational modifications in proteins specifically and at a single molecule resolution (40). Rudhira knockout cells showed significantly increased SMAD2/3 ubiquitination *in vivo* (marked by an increase in the number of PLA dots), both in the presence or absence of exogenous TGF β (Fig. 7C). This indicates that Rudhira promotes TGFβ pathway activation by negatively regulating inhibitors and thereby checking SMAD2/3 ubiquitination.

## Discussion

A balance of signal sensing and response is essential to maintain homeostasis and depends on the nature of the ECM and the responsiveness of the cytoskeleton, two very important parameters in determining cell phenotype during development and disease (29, 41, 42). The cytoskeletal response to signals would determine whether a cell can change its shape, divide or adhere. The cytoskeleton affects nuclear stability, which impinges on chromosome scaffolds and ultimately on gene expression (43). The generic role of the cytoskeleton in cell phenotype is acknowledged and even obvious, but there are only a few examples linking its organization to primary determinants of cell fate, as few tissue-specific cytoskeletal components are known. Context-specific cellular responses are likely to depend on the presence of such components, which use ubiquitous elements to shape context-dependent outputs. We therefore chose to analyse the function of Rudhira, a cytoskeletal protein expressed predominantly in the vasculature.

We report here that global or endothelial-specific deletion of *rudhira* resulted in mid-gestation lethality with severe defects in cardiovascular patterning and tissue morphogenesis. Based on a transcriptome analysis we identified key steps in blood vessel remodeling such as cell adhesion, migration and modulation of extracellular matrix components that are regulated by the action of Rudhira. We show the essential function of Rudhira in directing endothelial cell migration during development and describe, for the first time, a regulatory network mediated by Rudhira with reference to its interacting partners at both binary (regulatory) as well as physical levels (Fig. 3I). We propose a model by which Rudhira/BCAS3, an endothelial cell cytoskeletal component, can regulate microtubule-mediated TGFβ signaling (Fig. 7D).

In ECs TGFβ signaling modulates the transcriptome in a manner that would promote EC migration during angiogenesis (17). Also, in the yolk sac, paracrine TGFβ- signaling regulates gene expression and ECM production, deposition and remodeling, required for assembly of robust vessels (18). However, the absence of Rudhira hampers TGFβ signaling, negatively affecting endothelial cell migration. Expectedly, *rudhira* null ECs show a disorganized ECM. Matrix degradation is known to support angiogenesis in multiple ways such as by physical removal of barriers to migration, release of sequestered growth factors or by exposing cryptic protein sequences that function in angiogenesis (28, 44). MMP activity is essential for releasing TGFβ for growth factor signaling, which promotes cytoskeletal remodeling seen in mesenchymal migration. In the absence of Rudhira at least some of these angiogenesis promoting events do not occur, resulting in a disorganized matrix, possibly not permissible for directional migration. Though gelatin degradation is reduced upon *rudhira* depletion, expression of the cognate MMPs (2, 9 etc.) is not altered. We see significant increase in *MMP10, MMP21, MMP25* and reduced level of *MMP28* transcripts suggesting that this should cause increased degradation. However, MMPs are also regulated by TGFβ signaling. The reduced matrix degradation seen in *rudhira* CKO yolk sacs is in concordance with reduced TGFβ signaling.

Cardiac defects are seen from early development in *rudhira* mutants, suggesting impaired circulation. From E8.5 in the mouse yolk sac, blood flow dictates vessel fusion and directional cell migration resulting in vascular remodeling (45). Hence cardiac defects seen in *rudhira* mutants could also contribute to the vascular remodeling abnormalities and aberrant TGFβ signaling.

Rudhira overexpression in endothelial cells promotes TGFβ pathway activation. Further, addition of TGFβ induces Rudhira expression. This, in addition to other observations, indicates a positive feedback of Rudhira on TGFβ signaling and *vice versa.* In the absence of Rudhira this feedback is lost and it affects expression of TGFβ targets and the endothelial cell transcriptome. The upstream sequence of *rudhira* bears binding sites for the SMAD family of transcription factors (Smad3, Smad4), Twist subfamily of class B bHLH transcription factors, Pax (Paired box factors), Brachyury and ELK1 (member of ETS oncogene family). Further, our analysis shows that Rudhira functions after receptor activation but before or at the level of SMAD2/3 activation. This suggests that Rudhira may function to regulate transcription in response to TGFβ activation, possibly by interaction with other cytoskeletal and signaling components.

Rudhira localization is cytoskeletal and is dynamic in migrating cells that change shape and adhesion properties. Microtubules regulate TGFβ signaling by binding to and preventing activation of SMADs (19). Thus our analysis provides a platform for testing cell type-specific cross talk between TGFβ signaling and the dynamic cytoskeleton in normal development as well as pathological situations such as tumor metastasis. An increase in MT and Rudhira co-localization upon TGFβ stimulation suggests that Rudhira might serve to sequester microtubules thus inhibiting MT and Smad2/3 association and promoting SMAD2/3 phosphorylation (Fig. 6D). While targeting the interaction of MT with Smads would allow regulation of the TGFβ pathway, the ubiquitous nature of these molecules is likely to result in widespread and possibly undesirable effects of therapeutic intervention. Rudhira being restricted to the vasculature could provide a suitable tissue-specific target to regulate pathological angiogenesis and TGFβ pathway activation.

The presence of cell adhesion assemblies is dependent on the underlying matrix and on whether the surface is rigid as in 2D cell culture or embedded within a 3D ECM (46). Elucidating mechanisms by which Rudhira governs cell migration will be crucial to understanding tumor cell invasion and metastasis. Indeed the upregulation of Rudhira in metastatic tumors underscores its importance (47). Interestingly, TGFβ cross talks with several other signaling pathways and is a drug target for multiple diseases. Since TGFβ is also a key inducer of the epithelial/endothelial to mesenchymal transition, it is likely that the cytoskeletal remodeling in these transitions is mediated by Rudhira. Further studies on the role of Rudhira in normal and tumor EMT will be informative.

Given the large number of signals and their varying levels that EC encounter, it is unlikely that they respond only to one or the same concentration all across the organism. A robust response over a range of signals is likely mediated by molecules that can crosstalk with a wide variety of processes. The cytoskeleton is ideally positioned for this role. EC respond to a variety of signals resulting in a limited repertoire of cytoskeletal changes, which in turn determine cell phenotype. The presence or absence of cell type-specific components such as Rudhira could provide active decisive control of EC behaviour in response to a varying milieu of signals. Our analysis indicates that in endothelial cells Rudhira remodels actin (23) and affects TGF-P meditated gene expression (this report), which in turn results in matrix degradation, making the ECM permissive to cell migration. Thus our study opens up new avenues for developing strategies to regulate the vascular pattern in development and disease. This could be generally applicable to various cell and tissue types and identification of new components awaits further investigation.

## Materials and Methods

### Generation and validation of *rudhira* knockout mice

All animals were maintained and experiments performed according to the guidelines of the animal ethics committees. *Rudhira* floxed mice (Accession No. CDB0664K: http://www2.clst.riken.jp/arg/mutant%20mice%20list.html) were generated as described, validated by genotyping and crossed to *Cre* mice to generate knockout mice (see Supporting Information and Fig. 1 and S1 Fig.).

### RT- PCR and qRT-PCR

RNA from E9.5 embryos was isolated using TRIzol reagent (Invitrogen). Reverse transcription was performed using 2 μg of DNase treated RNA and Superscript II (Invitrogen, Carlsbad, CA) according to manufacturer’s instructions. Quantitative RT–PCR (qRT-PCR) was carried out using EvaGreen (BIO-RAD, CA) in Biorad-CFX96 Thermal cycler (BIO-RAD, CA). Primers used are provided in S7 Table.

### Immunostaining, Immunohistochemistry, Microscopy and analysis

Embryos were dissected at desired stages between E7.5 to E11.5, fixed in 4% paraformaldehyde and processed for cryosectioning or immunostaining using standard procedures (48). Primary antibodies used were against Rudhira (23), PECAM1, CD34, Flk1, Ly76 (BD Biosciences), Brachyury (Santa Cruz Biotechnology), laminin, fibronectin, Smooth Muscle Actin, acetylated tubulin, α-tubulin (Sigma Chemical Co. USA), SMAD2/3, pSMAD2 (Cell Signaling Technologies, USA). Secondary antibodies were coupled to Alexa-Fluor 488 or Alexa-Fluor 568 or Alexa-Fluor 633 (Molecular Probes). Cryosections were stained with haematoxylin and eosin using standard protocols. Images were acquired using a stereo zoom (SZX12 Olympus) or inverted (IX70, Olympus) microscope, confocal microscopes (LSM 510 Meta and LSM 700, Zeiss) and a motorized inverted microscope with fluorescence attachment (IX81, Olympus). For details, see Supporting Information. Co-localization analysis was done using colocalization plugin in ImageJ (NIH, USA).

### TGFβ induction and analysis

Live E10.5 and E11.5 CKO yolk sacs cut into two pieces or embryo-derived cells were washed in PBS and induced with 0 ng/ml (uninduced) or 10 ng/ml of TGFβ in DMEM for 2 h and then fixed and stained for pSMAD2. At least two each of the control and CKO E10.5 yolk sacs were taken for analysis. Non-silencing control (NS) or *rudhira* knockdown (KD) ECs were induced with 0 or 10 ng/ml of TGFβ in DMEM for 2 h and analysed by immunofluorescence or western blotting for SMAD2/3 and pSMAD2. Non-silencing control (NS) or *rudhira* knockdown (KD) ECs were induced with 0 or 10 ng/ml of TGFβ in DMEM for 48 h and analysed by qPCR for indicated genes or immunofluorescence for SMAD2/3.

### Western blot analysis and immuno-precipitation

50 μg lysate from control or knockdown cell lines of SVEC was used for Western blot analysis by standard protocols. Primary antibodies used were: SMAD2/3, pSMAD2 (Cell Signaling Technologies, USA), GAPDH, acetylated tubulin, α-tubulin (Sigma Chemical Co., USA), Ubiquitin (Clone FK2, Biomol) and BCAS3 (Bethyl Labs, USA). HRP conjugated secondary antibodies against appropriate species were used and signal developed by using Clarity Western ECL substrate (Biorad, USA). Western blot intensities were normalised to GAPDH and quantification was carried out using ImageJ. For immuno-precipitation studies, Protein G sepharose beads coated with SMAD2/3 antibody were incubated with cell lysates for 4 hours, clarified by centrifugation and extensively washed. Equal volumes of sample were loaded and resolved by SDS polyacrylamide gel and further taken for western blot analysis.

### *In situ* PLA reaction (Duolink assay)

*In situ* PLA reaction was performed on yolk sac primary cells. The cells were cultured, fixed, permeabilised and stained with primary antibodies for SMAD2/3 and Ubiquitin as mentioned earlier. Thereafter, the protocol for PLA as recommended by manufacturer (Duolink, USA) was followed. Post PLA, nuclei were counterstained with DAPI.

### *In situ* zymography

Unfixed embryos or yolk sacs were overlaid with tissue freezing medium, snap frozen in liquid nitrogen, cryosectioned at 10μm, collected onto slides and overlaid with zymography solution [20μg/ml DQ gelatin (Life Technologies, USA) in 50mMTris-HCl, 150mMNaCl, 5mM CaCh]. The slides were incubated at 37°C in the dark in a moist chamber for 15 min (yolk sac) or 2 h (embryos), then rinsed with ultrapure water (Milli-Q, Millipore) and fixed with 4% paraformaldehyde. Multiple sections from at least four embryos per genotype were analysed.

### Endothelial cell culture, adhesion and migration assays

E9.5 embryos or yolk sacs were washed in phosphate buffered saline (PBS), minced and dissociated in 0.2% collagenase type IV (GIBCO/BRL) at 37μC for 5 minutes, washed in culture medium, pelleted, resuspended in culture medium (DMEM, 20% fetal calf serum, 1X Glutamax, 1X antibiotics and 50 μg/ml Endothelial Cell Growth Supplement (ECGS) (Sigma Chemical Co., USA) and plated onto 0.1% gelatin coated dishes. Confluent monolayers were incubated with 5μg/ml DiI-Ac-LDL (Invitrogen) for 4 hours to mark endothelial cells, then scratched and monitored for cell migration in real time by video microscopy (see Supporting Information). For assessing effects on cell migration on different substrates, SVEC plated on desired matrix components were allowed to adhere and form a monolayer for 24 h before wounding. For assaying the effect of TGFβ on migration, 5ng/ml TGFβ was added to cultures 6 h before scratching and continuously provided up to 12h post-wounding. Wound width at 0 h and 12 h post-wounding was measured and quantified as described before (23).

### Microarray analysis

Stage-matched E9.5 embryos of control and ubiquitous knockout littermates were used for microarray analysis as detailed in Supporting Information.

### Quantification and Statistical analyses

Quantification and statistical analysis are described in Supporting Information.

## Acknowledgements

We thank staff at the Jackson Laboratories, USA for inputs on mouse breeding and maintenance; Developmental Studies Hybridoma Bank, University of Iowa, USA for some antibodies; JNCASR Imaging facility, NCBS Central Imaging and Flow Facility, JNCASR Animal Facility, NCBS Animal facility for access nd Inamdar laboratory members for fruitful discussions.

## Author Contributions

M.S.I. conceived of the project and directed the work. M.S.I., R.S., D.J., J.C.P., M.J., G.B., P.B. designed and performed animal experiments, cell biology, and imaging. R.S., M.V., M.S.I., D.J. analyzed transcriptome data. M.S.I. designed floxed mice and T.A., and H.K generated floxed mice. M.S.I., R.S., M.V., K.V.R. wrote the manuscript. All authors reviewed and made comments on the manuscript.

